# The interplay between lncRNAs, RNA-binding proteins and viral genome during SARS-CoV-2 infection reveals strong connections with regulatory events involved in RNA metabolism and immune response

**DOI:** 10.1101/2022.03.26.485903

**Authors:** Francisco J. Enguita, Ana Lúcia Leitão, J. Tyson McDonald, Viktorija Zaksas, Saswati Das, Diego Galeano, Deanne Taylor, Eve Syrkin Wurtele, Amanda Saravia-Butler, Stephen B. Baylin, Robert Meller, D. Marshall Porterfield, Douglas C. Wallace, Jonathan C. Schisler, Christopher E. Mason, Afshin Beheshti

## Abstract

Viral infections are complex processes based on an intricate network of molecular interactions. The infectious agent hijacks components of the cellular machinery for its profit, circumventing the natural defense mechanisms triggered by the infected cell. The successful completion of the replicative viral cycle within a cell depends on the function of viral components versus the cellular defenses. Non-coding RNAs (ncRNAs) are important cellular modulators, either promoting or preventing the progression of viral infections. Among these ncRNAs, the long non-coding RNA (lncRNA) family is especially relevant due to their intrinsic functional properties and ubiquitous biological roles. Specific lncRNAs have been recently characterized as modulators of the cellular response during infection of human host cells by single stranded RNA viruses. However, the role of host lncRNAs in the infection by human RNA coronaviruses such as SARS-CoV-2 remains uncharacterized. In the present work, we have performed a transcriptomic study of a cohort of patients with different SARS-CoV-2 viral load. Our results revealed the existence of a SARS-CoV-2 infection-dependent pattern of transcriptional up-regulation in which specific lncRNAs are an integral component. To determine the role of these lncRNAs, we performed a functional correlation analysis complemented with the study of the validated interactions between lncRNAs and RNA-binding proteins (RBPs). This combination of *in silico* functional association studies and experimental evidence allowed us to identify a lncRNA signature composed of six elements - NRIR, BISPR, MIR155HG, FMR1-IT1, USP30-AS1, and U62317.2 - associated with the regulation of SARS-CoV-2 infection. We propose a competition mechanism between the viral RNA genome and the regulatory lncRNAs in the sequestering of specific RBPs that modulates the interferon response and the regulation of RNA surveillance by nonsense-mediated decay (NMD).

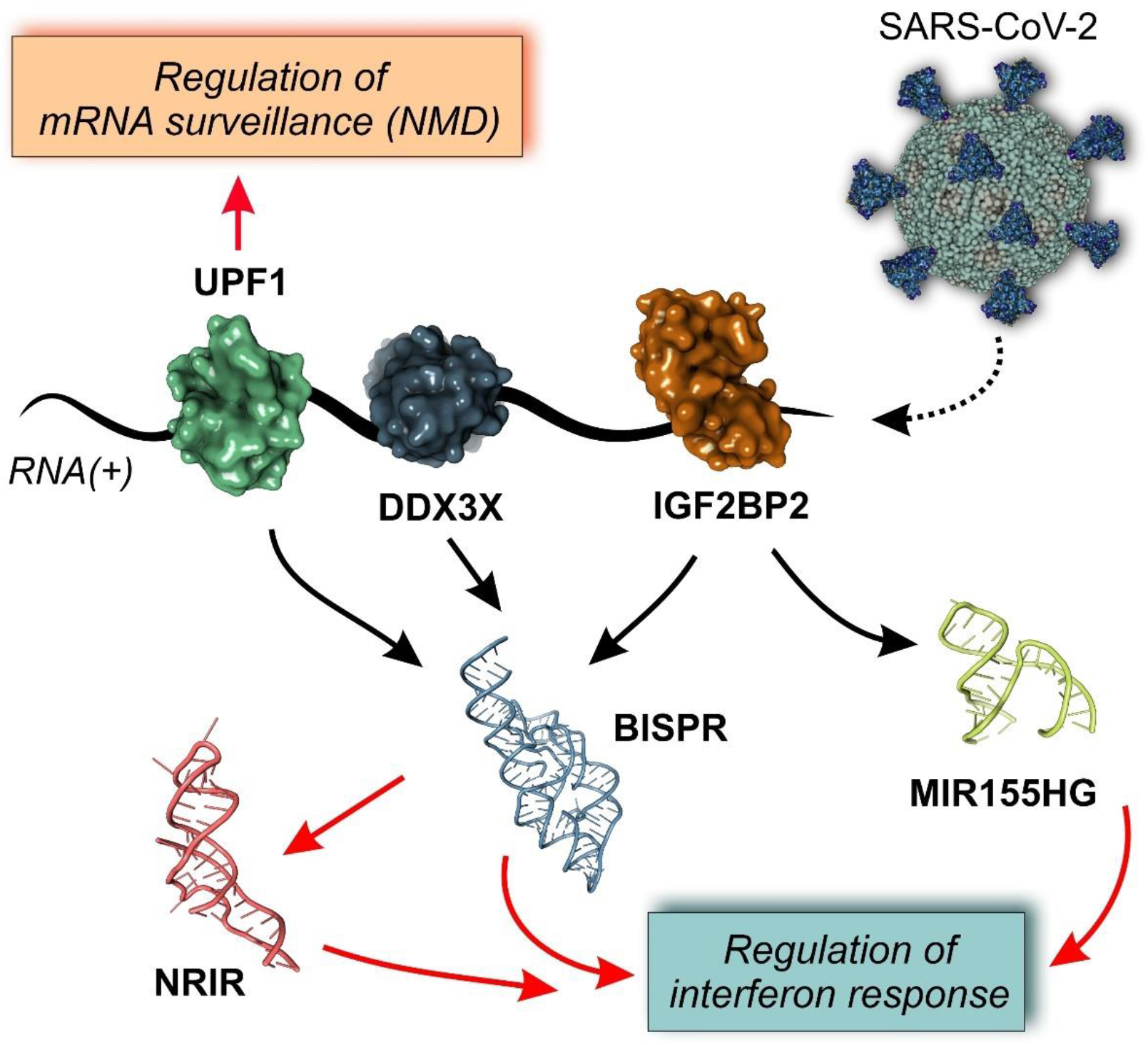

**Graphical abstract:** Model of interactions among lncRNA and cognate RNA-binding proteins in SARS-CoV-2 infection. According to our model, the viral genome can establish direct interactions with three core proteins (DDX3X, UPF1 and IGF2BP2) involved in mRNA metabolism and regulation of the interferon response, which are also components of a SARS-CoV-2 lncRNA-centered regulatory network. The competition between viral RNA and lncRNAs could act as a counteracting factor for the normal function of homeostatic lncRNA-centered regulatory networks, contributing to viral progression and replication. Black arrows depict physical interactions between network components; red arrows represent functional relationships.

## 1. INTRODUCTION

Pervasive transcription of the human genome generates a wide range of regulatory RNA molecules that control the flow of genetic information originated from the cell nucleus. Among these regulatory RNAs, long non-coding RNAs (lncRNAs), defined as those non-coding RNAs (ncRNAs) with sizes larger than 200 nucleotides and originated from specialized transcriptional units, are a very diverse class. These genes typically harbor their own promoters and regulatory sequences, many undergoing splicing and post-transcriptional modifications [1]. According to a recent update of the GENCODE database, the estimated number of lncRNA genes in the human genome is now over 18,000, a comparable number to the protein coding genes (around 20,000) [2]. LncRNA transcriptional units typically generate structured RNA molecules with regulatory functions that modulate the genomic output at different levels, including acting as scaffolds of high-molecular weight complexes as well as interacting with other biomolecules such as DNA, RNAs, and proteins [3-5]. LncRNAs have been found to have cell-state specific functions and have modulating effects on protein-coding gene transcription [6]. Transcriptomic analysis driven by next-generation sequencing applications has unveiled functional relationships between lncRNA and the pathophysiology of metabolic diseases, cancer, and infections [7-9]. The existence of a pathology is often accompanied by a dysregulation of lncRNA expression that could represent a secondary event associated with the disease or as a driving factor of the condition [10].

Viral infections are extreme cases of the interaction between two organisms in which the infectious agent strictly depends on the molecular and metabolic machinery of the infected cell to complete its replication and proliferation cycle. During the hijacking of the host cellular machinery by the virus, key molecular interactions between viral components and cellular structures are established. These interactions are responsible for the reorganization of cellular membranes to facilitate virus entry, modulation of cellular metabolism, and evasion of specific defense mechanisms [11]. Most of the knowledge about cellular and viral molecular players during infection is in the protein realm, represented by the characterization of viral-encoding polypeptides that are responsible for the progression of the infection or immune evasion, and their cellular cognate targets. However, the relevance of cellular and viral RNAs as relevant players within the context of an infection must be considered [12].

The roles of lncRNAs as mediators or drivers of viral infections were first unveiled in the last decade [13]. LncRNA mediators have been shown to play key roles in the regulation of the immune and inflammatory response against viral infections [14, 15]. These well-described examples define the regulatory role of individual lncRNAs during RNA viral infections [16-18]. For instance, a leading cause of viral gastroenteritis from the human norovirus can induce a strong lncRNA-based response in the host cells that is related to the regulation of the interferon response [19]. Strains of human hepatitis C virus (HCV) associated with long-term persistence downregulate the expression of lncPINT (p53-induced transcript long non-coding RNA) as a mechanism for circumventing the interferon defense mechanism and evading the innate immune response [20]. Following a similar strategy, the recently characterized lncRNA AP000253, provides a mechanism by which hepatitis B virus can remain occult for prolonged times within the host [21]. In many of these examples, results obtained from experimental models linked the lncRNA mediators of infection with a complex network of RNA-binding proteins (RBPs) [20, 22].

SARS-CoV-2, a respiratory RNA(+) virus with a rapid transmission pattern, was responsible for the global pandemic that started in late 2019. SARS-CoV-2 is a virus belonging to the *coronaviridae* family that enters the cell by specific interactions with the host ACE2 receptor [23, 24]. After internalization, cell infection is characterized by a dysregulated gene and protein expression pattern that includes an up-regulation of genes involved in the interferon response and interleukin production [25, 26]. If the virus evades host cell defenses, the replication of the genetic material is enabled by a multimeric RNA-dependent RNA polymerase. The RNA genome is translated into a polypeptide that is matured by proteolytic specific digestion with two viral proteases, the main protease (MPro) and the papain-like protease (PLPro) [27]. Whole virions are assembled and secreted by a pathway that involves the participation of the endoplasmic reticulum and Golgi complex [11, 12]. In severe cases, SARS-CoV-2 infected patients showed a striking pattern of acute inflammatory responses that has been related to the uncontrolled production of cytokines and designated as “cytokine storm” [28, 29].

Genomic SARS-CoV-2 RNA and SARS-CoV-2 RNA transcripts interact with specific proteins modulating cellular responses to the infection, as revealed by high-throughput proteomic analysis [25, 30]. The multiple interactions between the viral genome/transcriptome and cellular proteins are a factor in promoting replication of the virus or, contrariwise, ensuring the success of the cell in preventing replication [31, 32]. Small non-coding RNAs (ncRNAs) have been described as regulatory factors in the virus-host interface [33]. However, the functions and roles of lncRNAs in the development and progression of SARS-CoV-2 infection remain uncharacterized. We determined the lncRNA dysregulation pattern induced by the SARS-CoV-2 infection and characterized the lncRNA-centered regulatory networks involving RBPs associated with RNA metabolism and interferon-mediated responses, by analysis of high-throughput transcriptomes of samples obtained from patients with and without SARS-CoV-2 infection. The detailed knowledge of the complex regulatory networks involving lncRNAs could open new perspectives for the design of targeted drugs to treat severe cases of SARS-CoV-2 infection.

## 2. MATERIAL AND METHODS

### 2.1. Data source and group stratification

The source data for this study was generated within the framework of COV-IRT consortium (www.cov-irt.org) and deposited at the Short Read Archive (SRA) database with the project reference PRJNA671371, corresponding to a previously published study [26]. The dataset includes a shotgun metatranscriptomic (total RNA-seq) for host and viral profiling of 735 clinical specimens obtained from patients at the Weill Medical College of Cornell University, New York, USA. Patients were stratified according to the SARS-CoV-2 levels determined by qRT-PCR experiments by simultaneously using primers to amplify the E (envelope protein) and S (spike protein) genes together with the proper internal controls as previously described [26]. Patients with a cycle threshold value (Ct) less than or equal to 18 were assigned to “high viral load”, a Ct between 18 and 24 were assigned to “medium viral load”, and a Ct between 24 and 40 were assigned to “low viral load” classes, with anything above a Ct of 40 classified as “negative” [26]. These last patients were also subdivided according to the presence of other viral respiratory infections different from Covid19 and having compatible symptoms.

### 2.2. Analysis of RNAseq data

Raw Illumina sequence reads obtained by a pair-end sequencing strategy, were filtered, and trimmed with Trimmomatic software [34]. Filtered sequence reads were dual-aligned with the reference SARS-CoV-2 genome from Wuhan (strain reference MN908947.3) and the human genome (genome build GRCh38 and GENCODE v33) using the STAR aligner [35]. The gene counts were indexed to the different families of coding and non-coding gene transcripts by the BioMart data portal [36]. Data was normalized using the variance-stabilizing transform (vst) in the DESeq2 package [37]. Differential gene expression between working groups was determined by the Limma/Voom algorithm implemented in the iGEAK data processing platform for RNAseq data [38]. Criteria for selection of significant differentially expressed genes included an adjusted p-value < 0.05, and logFc < -1.0 or logFc > 1.0. All the gene expression data is publicly available at the Weill Cornell Medicine COVID-19 Genes Portal, an interactive repository for mining the human gene expression changes in the data from this study (covidgenes.weill.cornell.edu).

### 2.3. Bioinformatic analysis of lncRNA-centered regulatory networks

The functional annotation of the group of selected lncRNAs whose expression was induced by SARS-CoV-2 infection was performed by the ncFANS 2.0 platform using the ncRNA-NET module [39]. Applying this module, we determined the co-expression network involving the selected lncRNAs and protein-coding genes using data extracted from healthy tissues and compiled in the Genotype-Tissue Expression (GTEx) portal [40]. The correlated coding genes were functionally grouped by GO-term analysis, pathway enrichment, and determination of molecular signatures by the ncRNA-NET module in ncFANS. In addition to the classical GO-term enrichment analysis, the redundant ontology terms were filtered by REVIGO software [41].

The lncRNA-centered regulatory networks established between lncRNAs and RNA-binding proteins were constructed by interrogating the ENCORI database for RNA interactomes [42]. Graphical analysis and representation of lncRNA-centered regulatory networks was performed by NAViGaTOR software [43]. Functional similarity of the selected overexpressed lncRNAs in SARS-CoV-2 infection was inferred by integrating heterogeneous network data with IHNLncSim algorithm [44]. This approach integrates information from experimentally validated data at three levels of functional association: miRNA-lncRNA, disease-based correlation and GTEx expression-based networks.

## 3. RESULTS

### 3.1. Host transcriptional shift induced by SARS-CoV-2 infection

To characterize the cellular response against SARS-CoV-2 infection, we performed a transcriptomic analysis from nasopharyngeal swabs collected from patients testing for SARS-CoV-2 virus. The patients were previously stratified according to the presence or absence of positive qPCR test, the existence of other respiratory pathogens different from SARS-CoV-2 and the different virus loading depending on the amplification Ct parameters as described in the Material and Methods section. The results, depicted in Figure 1a, characterize the transcriptional dysregulation in the host cells associated with infections by SARS-CoV-2 and other respiratory viruses. In SARS-CoV-2 patients, increased viral load resulted in an increment of the number of upregulated transcripts (Figure 1b, 1c). Globally, the number of transcripts in infected patients with a logFC > 1 compared to uninfected control patients increased from 52 to 891 from low to high SARS-CoV-2 viral loads. Analyzing the different families of transcripts, high viral load SARS-CoV-2 infected patients together with those infected with other respiratory viruses showed a preferential upregulation pattern, where the coding RNAs were more abundant. Moreover, the patients with higher SARS-CoV-2 loads also showed greater proportion of upregulated transcripts represented by lncRNAs (Figure 1b).

**Figure 1:**
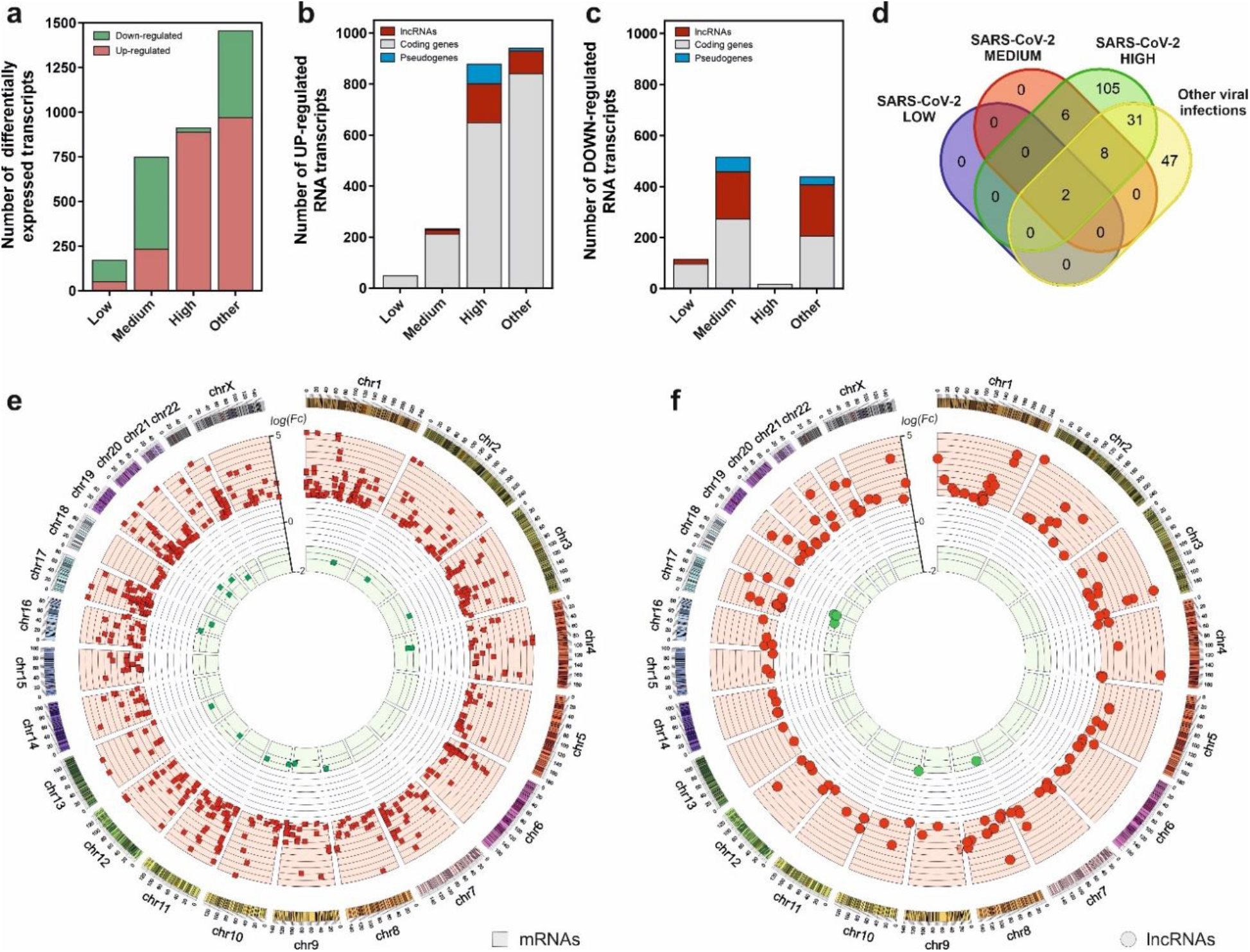
SARS-CoV-2 infection is characterized by a gene expression pattern enriched in up-regulated mRNA and lncRNA transcripts that can be correlated with the viral load observed in patients. **a**, number of differentially expressed transcripts observed in patients with different SARS-CoV-2 viral loads (Low, Medium and High) and those infected with different respiratory viruses (Other) in comparison with the uninfected patients; **b**, number of the different families of up-regulated transcripts in SARS-CoV-2 patients and infected with other respiratory viruses in comparison with the control group; **c**, number of the different families of down-regulated transcripts in SARS-CoV-2 patients and infected with other respiratory viruses in comparison with the control group; **d**, Venn diagram representing the number of up-regulated lncRNA transcripts observed in each group of study referred to the uninfected control group; **e**, CIRCOS plot [45] showing the genomic location and fold changes of the differentially expressed coding transcripts in the group of SARS-CoV-2 patients infected with higher viral loads in comparison with the uninfected controls (red squares, up-regulated mRNAs; green squares, down-regulated mRNAs); **f**, CIRCOS plot [45] depicting the genomic locations and fold changes of the differentially expressed lncRNA transcripts in the group of SARS-CoV-2 patients infected with higher viral loads in comparison with the uninfected controls (red circles, up-regulated lncRNAs; green circles, down-regulated lncRNAs).

Interestingly, from the 152 upregulated lncRNAs in high viral load samples, only 2 are common to all the analyzed infections. In SARS-CoV-2 infected patients, 105 upregulated lncRNAs were exclusive to the higher-level infections (Figure 1d). Positional gene enrichment analysis [46] of the upregulated lncRNA loci in high level SARS-CoV-2 infection showed two genomic regions enriched in overexpressed transcriptional units in response to the virus, comprising chr1: 148290889-155324176 and chr17: 32127595-62552121. The remaining overexpressed lncRNAs and coding mRNAs were evenly distributed across the different chromosomal loci with no evident spatial enrichment pattern (Figure 1e and 1f). The complete list of differentially expressed genes in all the comparisons is available as supplementary table (Suppl. Table 1).

### 3.2. Functional analysis of upregulated lncRNAs in SARS-CoV-2 infection

Prediction of lncRNA functions using the principles of systems biology is a challenging task due to the lack of supporting experimental evidence and the complexity of interactions established among lncRNAs and other functional players. Among the computer-based strategies available, we selected ncFANs 2.0 as a functional classifier [39]. The ncFANs-NET module was used to predict the functions of the upregulated lncRNAs in high-viral load infections by using the “guilty by association” approach. A correlation network between the differentially overexpressed lncRNAs and coding genes was constructed by ncFANs using data extracted from GTEx project database [47] and enrichment analyzed using terms from the Gene Ontology (GO) [48, 49] and KEGG databases [50]. The results of the functional analysis of the resultant co-expression network by GO-term enrichment with redundant term filtering, pathway analysis and molecular signature determination are depicted in Figure 2. GO-term enrichment within the category of molecular function resulted in the selection of terms related with cell-to-cell communication, and the general processes of lymphocyte activation and cytokine production (Figure 2a). The KEGG-pathway enrichment analysis resulted in a list of pathways also related with cytokine response and regulation, T-cell signaling and infections by viruses, bacteria, *Trypanosoma* and *Apicomplexa* parasites (Figure 2b), suggesting the common lncRNA-related regulatory responses exerted by the host cells against different infectious agents. Interestingly, the analysis of the molecular signatures in the regulatory lncRNA network revealed the existence of genes related with the interleukin signaling pathways, the interferon gamma response and the epithelial to mesenchymal transition phenomena, as more significant functions (Figure 2c).

**Figure 2:**
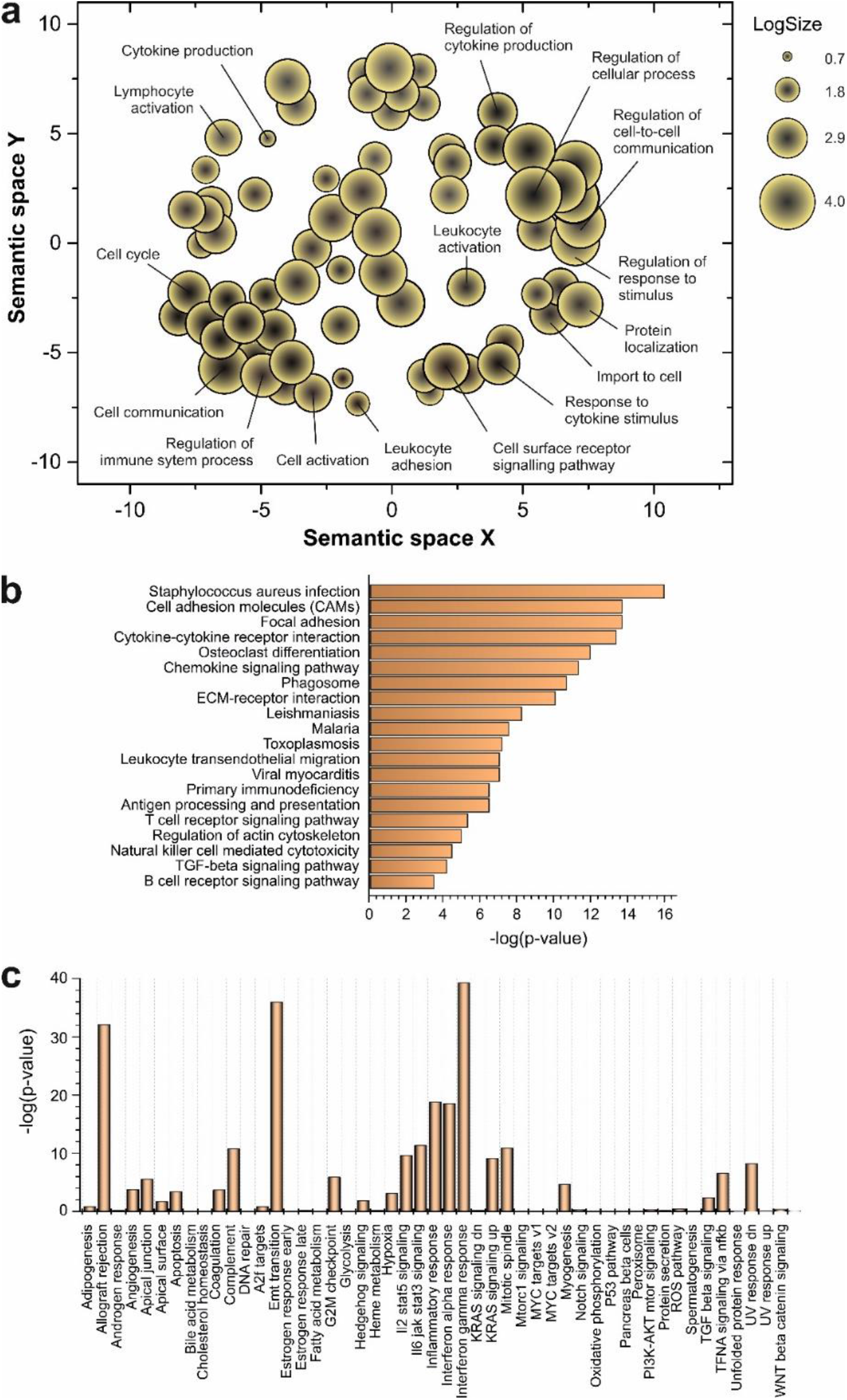
Functional prediction analysis by ncFANs 2.0 algorithm [39] of the upregulated lncRNAs observed in SARS-CoV-2 patients with high viral loads. **a**, GO-term enrichment analysis performed with ncFANs and filtered by removal of the redundant terms with REVIGO [41]. The filtered GO-terms are classified according to their two-dimensional arbitrary semantic space and represented by symbols with dimensions proportional to the LogSize, showing the most relevant GO-terms for the context of viral infections. **b**, pathway enrichment analysis by ncFANs using the KEGG database. **c**, molecular signature analysis by ncFANs using the MSigDB database.

### 3.3. Expression of lncRNAs involved in the regulation of interferon response is correlated with SARS-CoV-2 viral load

Considering the group of sample patients with higher SARS-CoV-2 viral loads, the top list of upregulated lncRNAs includes important non-coding transcripts previously described as regulators of the interferon-mediated immune responses. The expression of these lncRNAs across the different groups of patients is depicted in Figure 3. NRIR, a driver of the interferon response [51], showed an upregulation pattern in patients infected with SARS-CoV-2 and other respiratory viruses (Fig. 3a). A similar pattern can be observed in BISPR (Fig. 3c), an interferon-induced lncRNA [52] and MIR155HG (Fig. 3e), a lncRNA related to cell proliferation and the regulation of innate immune response against specific viral infections [53, 54]. The USP30-AS1 lncRNA is an antisense transcript to the USP30 gene that has been implicated in mitochondrial quality control in some cancers and also in the progression of virus-induced cancers such as malignant cervical tumors [55, 56], and is mainly upregulated in those SARS-CoV-2 patients with higher viral loads. Interestingly, the careful analysis of COVIDOME database [57] allowed us to determine that NRIR and BISPR lncRNAs are also upregulated in the blood of Covid19 patients (Fig. 3g and 3h). LINC02068 and AL512306.2 also showed an upregulation pattern which is dependent on the SARS-CoV-2 viral load in our patient cohort (Fig. 3b and 3f) that can be also observed in the COVIDOME dataset for AL512306.2 transcript (Fig. 3j).

**Figure 3:**
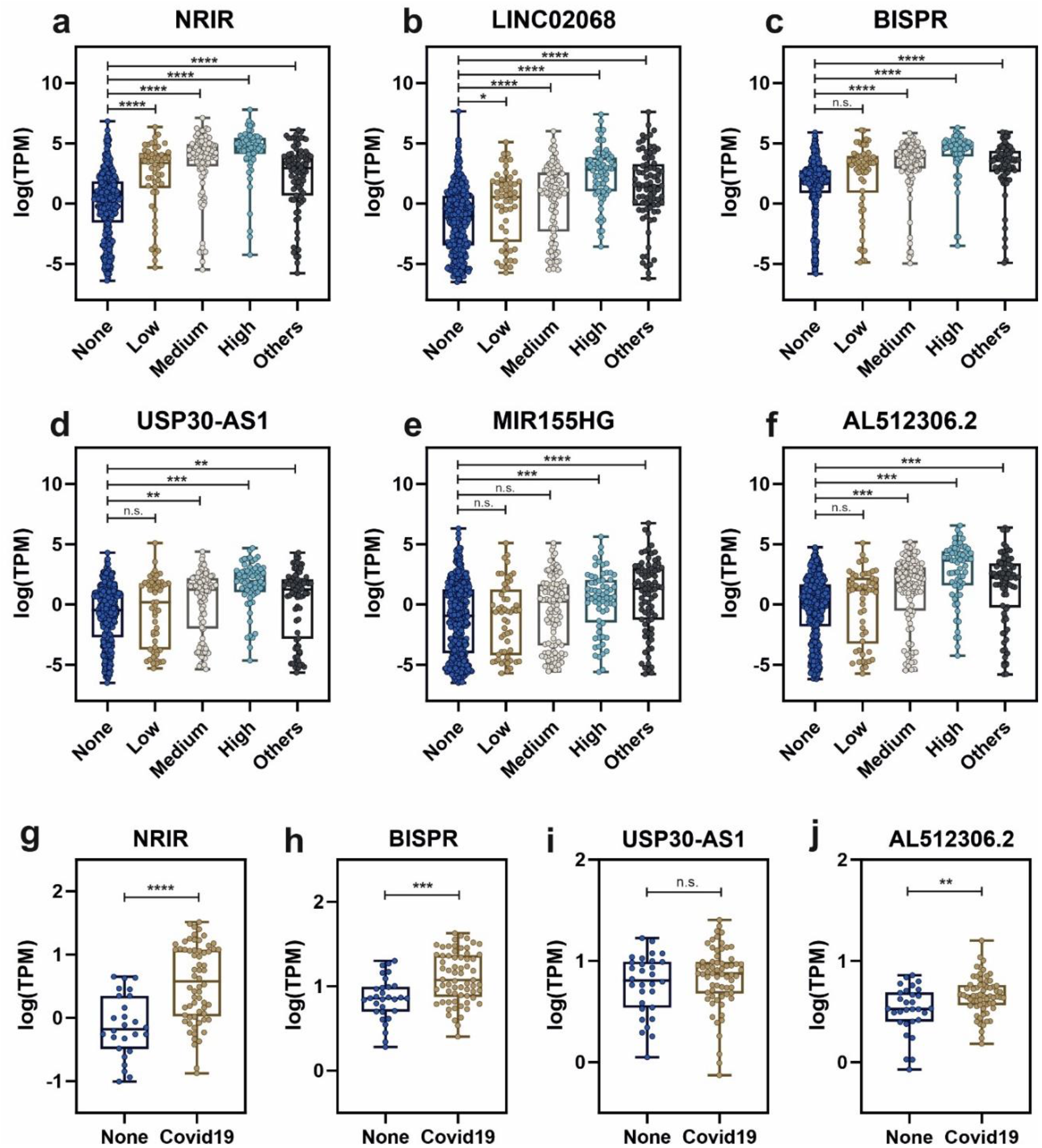
Expression levels of selected lncRNAs quantified by next-generation sequencing in nasal swabs from the working group of patients, and whole blood, obtained from the COVIDOME project database [57]. LncRNA expression in nasal swabs distributed by groups of patients: **a**, NRIR; **b**, LINC02068; **c**, BISPR; **d**, USP30-AS1; **e**, MIR155HG and **f**, AL512306.2. LncRNA levels in whole blood in SARS-CoV-2 patients (**Covid19**) and non-infected controls (**None**): **g**, NRIR; **h**, BISPR; **I**, USP30-AS1 and **j**, AL512306.2. Statistical comparisons between sample groups were made by one-way ANOVA in the case of samples from nasal swabs and by the Student’s t-test in the data from the COVIDOME project (****, p-value < 0.0001; ***, p-value < 0.001; **, p-value < 0.01; *, p-value < 0.05 and n.s, non-significant).

The results, depicted in Figure 4, show the Spearman’s correlation analysis of the 90 most upregulated lncRNAs in high-load SARS-CoV-2 infections across all the studied samples. Correlation matrix (Fig. 4a) clearly demonstrates the presence of clusters of lncRNAs with high correlation values that could respond to the existence of regulatory blocks defined by these lncRNAs. Next-generation sequencing data allowed us to detect SARS-CoV-2 transcripts in the patients’ samples, the mRNA encoding for the Spike protein being the most abundant. Analyzing the correlations of S-protein mRNA with the overexpressed lncRNAs, we found moderate positive correlation with NRIR (Fig. 4b) and BISPR (Fig. 4c) lncRNAs and low positive correlation with MIR155HG (Fig. 4d).

**Figure 4:**
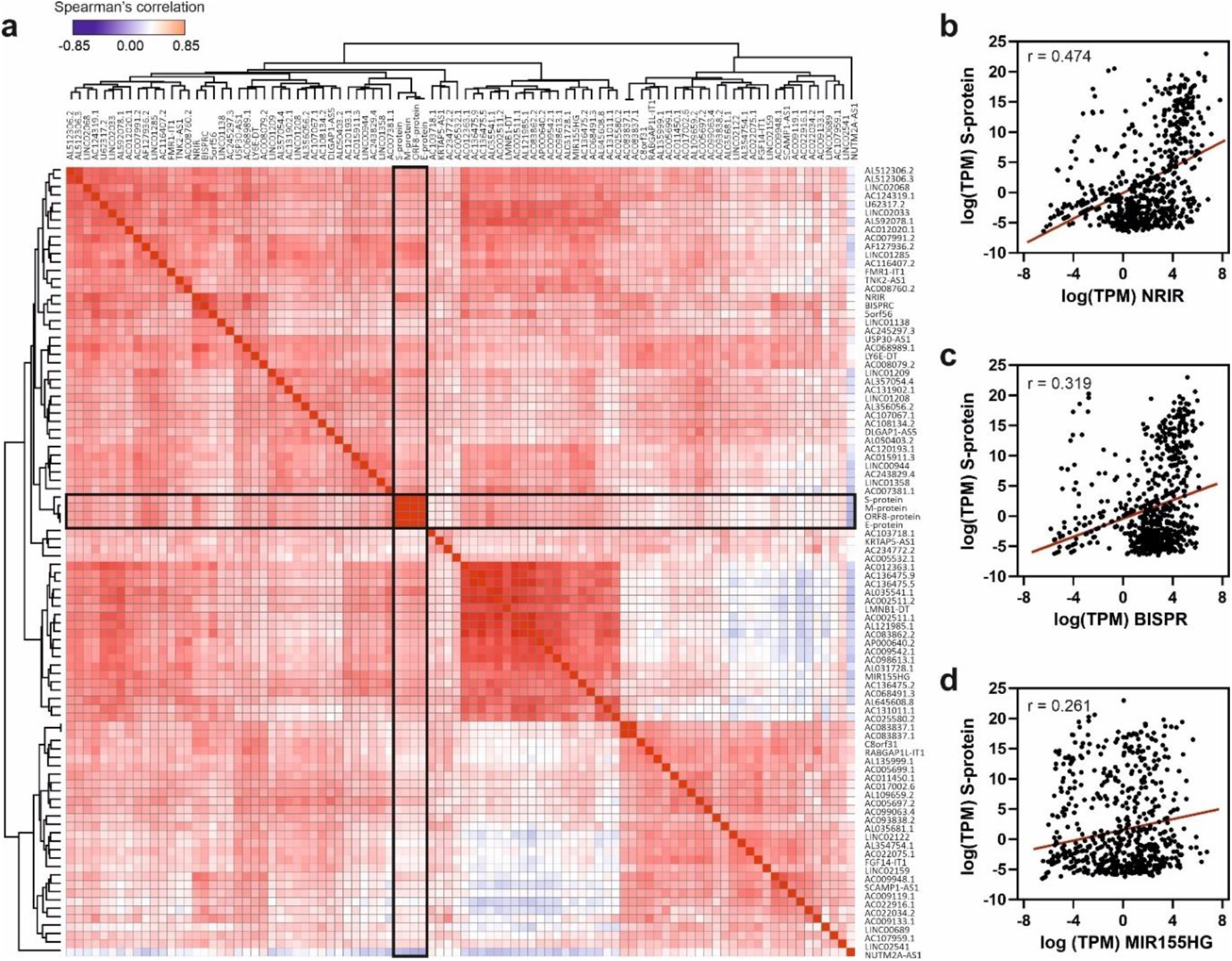
Spearman’s correlation analysis of the top 90 upregulated lncRNAs and the viral gene transcripts in the cohort of analyzed samples. **a**, Hierarchical clustered Spearman’s correlation matrix for the overexpressed lncRNAs and the detected SARS-CoV-2 transcripts across all the samples analyzed by the BioCPR software [63]. The SARS-CoV-2 mRNA transcripts are highlighted within boxes; **b**, correlation analysis between NRIR lncRNA and S-protein transcript; **c**, correlation analysis between BISPR lncRNA and S-protein transcript; and **d**, correlation analysis between MIR155HG lncRNA and S-protein transcript. The correlation coefficients showed in panels b, c and d correspond to the Spearman analysis and are significant in all cases with p-values < 0.0001.

### 3.4. Functional links between RNA-binding proteins and lncRNAs in SARS-CoV-2 infection

LncRNA function is exerted by their interaction with other biomolecules, namely DNA, other RNAs and proteins, and by the establishment of high-order molecular complexes where they can act as scaffolds and active regulatory players. To infer the possible role of the upregulated lncRNAs during SARS-CoV-2 infection we analyzed their interactions with RBPs to construct a network of functional relationships. We interrogated the ENCORI database of validated protein-RNA interactions [42] using the group of upregulated lncRNAs detected in patients with high SARS-CoV-2 loads and performed a functional enrichment analysis (Figure 5). The GO-term analysis of biological processes of the RBPs interacting with the upregulated lncRNAs showed an enrichment pattern of events related with RNA splicing, RNA metabolism and regulation of mRNA processing (Fig. 5a). On the other hand, a pathway enrichment analysis based on KEGG database of the selected RBPs demonstrated significant enrichment in splicing, mRNA surveillance pathways, RNA transport and, interestingly, some already described viral-related processes as viral-induced carcinogenesis and viral endocarditis (Fig. 5b).

**Figure 5:**
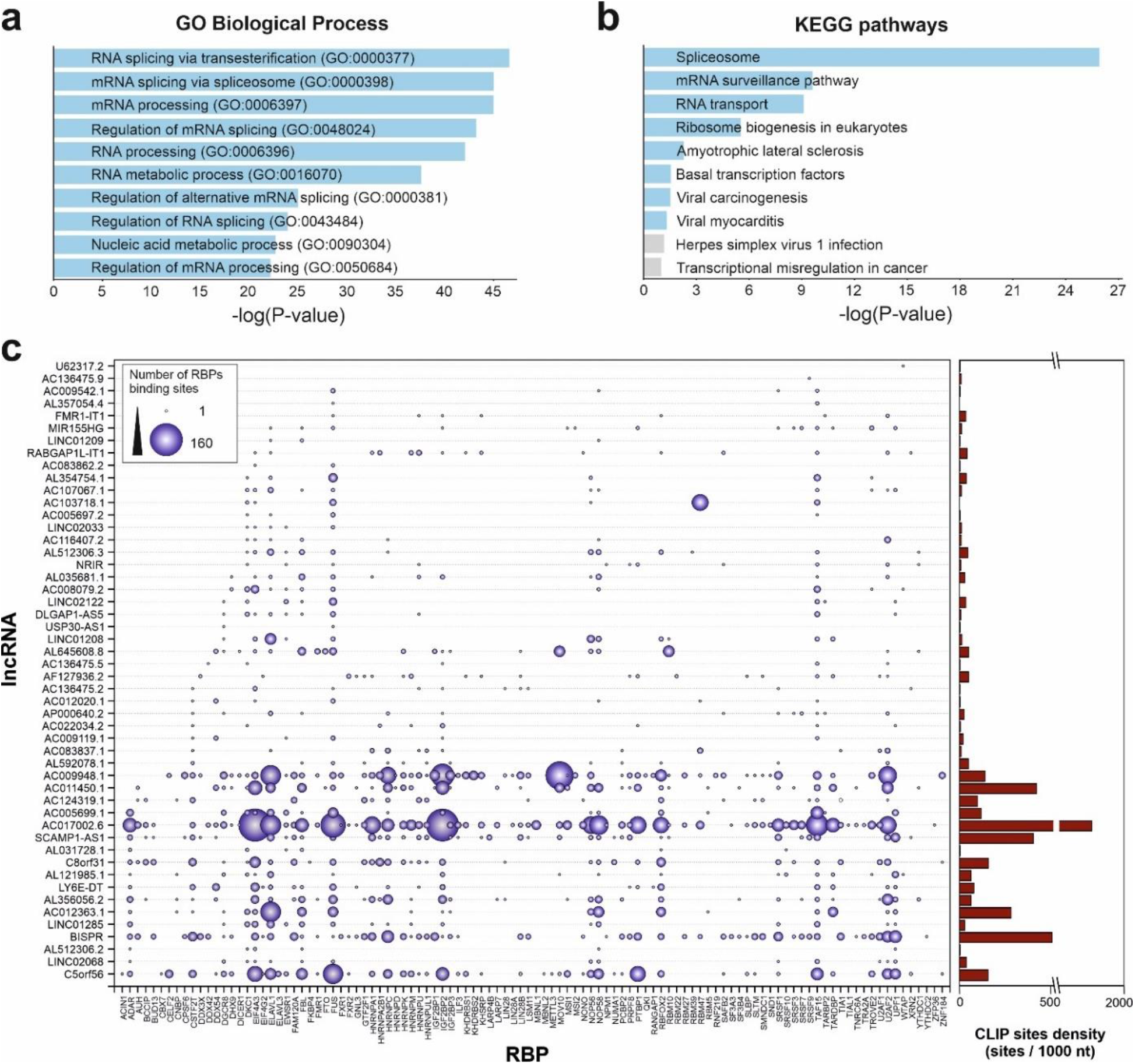
Functional analysis of the 50 top up-regulated lncRNAs by SARS-CoV-2 infection considering their validated interactions with RNA-binding proteins retrieved from ENCORI database [42]. **a**, GO-term enrichment analysis for biological processes of the RNA-binding proteins that interact with the selected overexpressed lncRNAs in SARS-CoV-2 patients with high viral loads; **b**, pathway enrichment analysis using the KEGG database and considering the RNA-binding proteins that interact with the selected overexpressed lncRNAs in SARS-CoV-2 patients with high viral loads; **c**, interaction map between RNA-binding proteins and the 50 top overexpressed lncRNAs in SARS-CoV-2 patients with high viral loads. The number of interactions is depicted as circles with a diameter proportional to the number of RNA-binding sites in each lncRNA. The right-hand side panel represents the density of RBP binding sites per 1000 nucleotides in each lncRNA as extracted from the ENCORI database.

To gain insights into the associations between RBPs and the Covid19-induced lncRNAs, we constructed a detailed interaction map by using the ENCORI data (Fig. 5c). This connection map includes information about the specific RBPs interacting with the upregulated lncRNAs together with the density of RBP-binding sites in each lncRNA determined by CLIP experiments and extracted from the ENCORI database [42]. Considering the global number of RNA-binding sites determined by CLIP experiments, the most represented RBPs comprised the EIF4A3 helicase, core of the exon-junction complex [58], the FUS transcriptional regulator involved in DNA repair, transcription and splicing [59], the TAF15 transcriptional regulator [60], the ELAVL1 regulator of RNA stability [61], and the IGF2BP2 protein, a previously known regulatory player that can interact with several ncRNAs including miRNAs and lncRNAs [62].

Additional overrepresented RBPs include splicing factors U2AF2, FBL and CSTF2T, and the nucleolar proteins NOP56 and NOP58. We could distinguish two groups of upregulated lncRNA transcripts, depending on the density of RBP-binding sites (Figure 5c). LncRNAs AC017002.6, AC107959.1 and BISPR showed a high-density of RBP-binding sites (>500 CLIP sites per 1000 nucleotides); this is compatible with their involvement in regulatory events related with the capture of RBPs by RNA sponging [64, 65]. On the other hand, lncRNAs as NRIR, MIR155HG, U62317.2, USP30-AS1 and TNK2-AS1 contain a reduced number of RBP-binding sites (<2 CLIP sites per 1000 nucleotides), thus are likely involved in the activity of lncRNA-centered regulatory complexes [51, 53].

### 3.5. Interplay of RNA-binding proteins between host and viral RNAs

Experimental evidence obtained in cell and animal model systems, described the existence of lncRNA-centered regulatory networks that modulate gene expression at different levels [66]. These regulatory networks are often composed of many lncRNAs that work in a coordinated manner to exert a regulatory action over a specific pathway [6]. In the contact of an external stimuli such as an infection, cells trigger a complex response where lncRNAs are important players [8, 21, 67]. Since we determined the existence of an upregulation lncRNA pattern after SARS-CoV-2 infection, we hypothesized about the existence of a coordinated lncRNA network that could regulate the cellular response to the virus.

To evaluate the existence of putative lncRNA-based regulatory modules in Covid19 response, we computed the similarity scores of all observed upregulated lncRNAs in patients with high viral loads using the IHNLncSim algorithm [44]. The results showed the presence of a group of 6 upregulated lncRNAs with significant similarity scores computed by two of the modules within IHNLncSim, the NONCODE-net and the lncRNA-Disease-net modules. The lncRNA signature includes NRIR, BISPR, MIR155HG, USP30-AS1, FMR1-IT1 and U62317.2 non-coding transcripts (Fig. 6a). NRIR and BISPR transcripts are virus-responsive lncRNAs involved in the regulation of the innate immune response and interferon signaling [52, 68]. MIR155HG has been also described as a regulator of the cellular response against influenza viruses [53] and recently characterized as upregulated in a cellular model of SARS-CoV-2 infection [69]. USP30-AS1 transcript is related to autophagy and mitochondrial quality control in the context of tumor progression [56, 70], whereas FMR1-IT1 and U62317.2 have no characterized functions.

**Figure 6:**
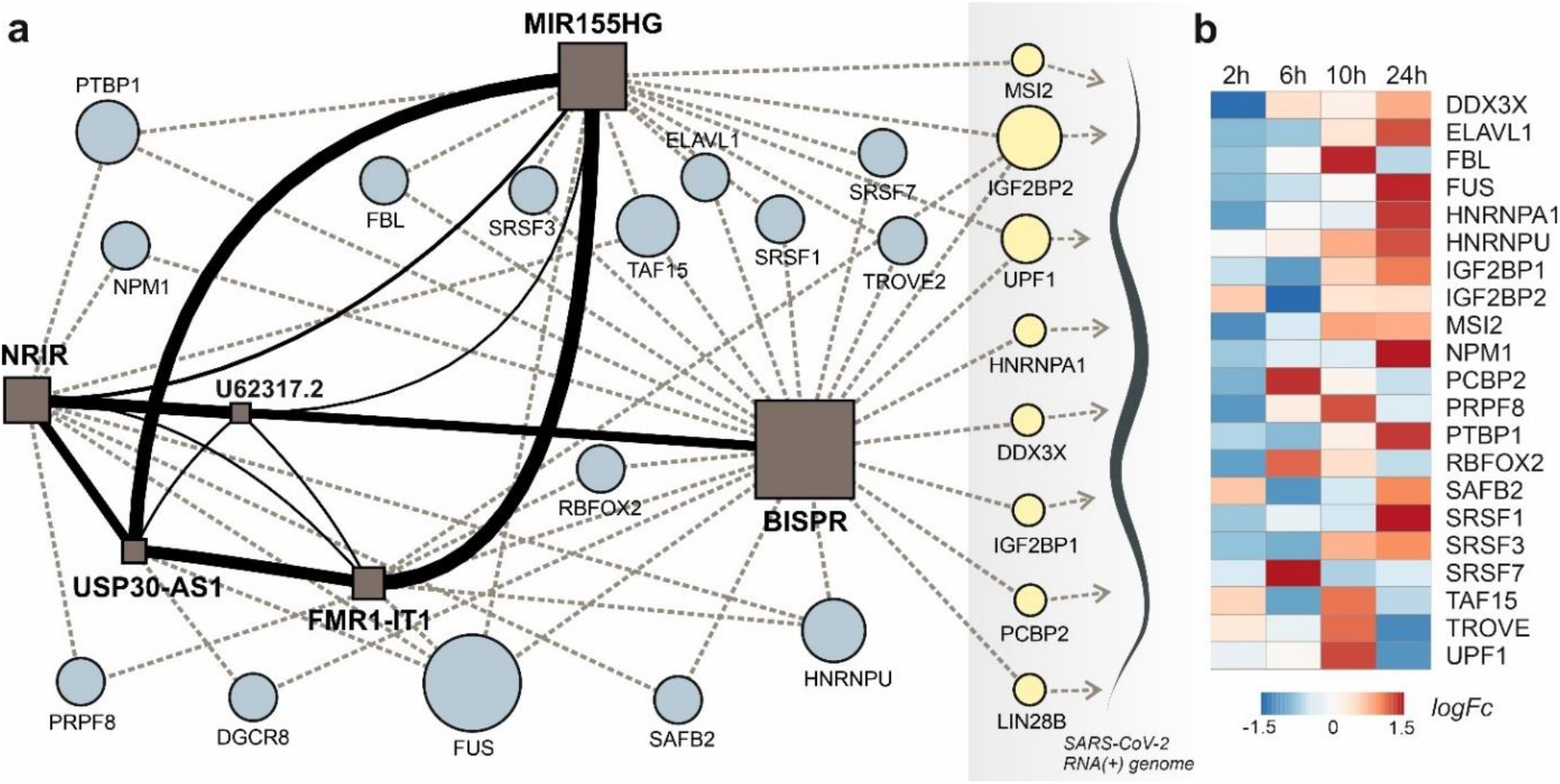
lncRNA-centered regulatory network established in SARS-CoV-2 infection involving upregulated lncRNAs, RNA-binding proteins and the viral genome. **a**, regulatory network built by heterogeneous network data analysis with IHNLncSim algorithm [44], the RNA-binding proteins extracted from ENCORI database [42] and the recently described interactions between host proteins and the viral genome [32]. Functional similarity among upregulated lncRNAs determined by IHNLncSim are represented by connecting continuous lines with thickness proportional to the value of the similarity coefficient value. Validated RNA-protein interactions from ENCORI database are represented by dashed grey lines. Characterized interactions between RNA-binding proteins and the SARS-CoV-2 genome are represented by dashed blue arrows. The size of the symbols representing lncRNAs (squares) and RNA-binding proteins (circles) are proportional to the number of established functional interactions. **b**, time course of protein expression from the selected RNA-binding proteins during SARS-CoV-2 in a cellular model, as described previously [25]. Expression data from quantitative proteomics were retrieved from the PRIDE partner repository database.

This lncRNA signature can be complemented with two additional layers: the RBP interaction network and the connections between these RBPs and the SARS-CoV-2 genome, recently characterized in cellular models of infection [31, 32]. Regarding the lncRNA-RBP interactions, except for BISPR lncRNA, all the members of the lncRNA signature belong to the group of transcripts with low RBP-binding site density as described in 3.4. Additionally, the analysis of the functional connections between lncRNAs and the virus genome allowed to select a group of RBPs involved in direct interactions with SARS-CoV-2 genome and the members of the lncRNA network simultaneously (Fig. 6a). This group of RBPs comprises two helicases (UPF1 and DDX3X), three generic RBPs (IGF2BP1, IGF2BP2 and LIN28B), a poly-C-binding protein (PCBP2), a ribonucleoprotein (HNRNPA1) and a translational inhibitor (MSI2). Taking advantage of the quantitative proteomics data already available in a cellular model of SARS-CoV-2 infection [25] we analyze the time course of the selected RBPs expression during Covid19 infection (Fig. 6b). During an infection time frame of 24h, the protein levels of the selected RBPs increased following the progression of the virus, showing maximum values at 10 or 24 hours of infection depending on the protein.

## 4. DISCUSSION

Viruses interact with cellular factors to complete their replicative cycles, avoiding the cell antiviral mechanisms. The dissection of the tangled network of interactions established during viral infections is essential not only to understand viral biology but also to develop targeted antiviral therapeutics. Among host factors regulating the cellular responses against viral infections, lncRNAs have been recently described as important players in host-viral interactions [71]. LncRNAs are a diverse family of ncRNAs with regulatory potential exerted mainly by their presence in functional complexes integrated by proteins and other RNA molecules [72].

Within the framework of the COVID-19 International Team (COV-IRT), we used an integrative approach to dissect the SARS-CoV-2 infection mechanisms and their physiological consequences. As part of our multidisciplinary research efforts, we studied the role of host lncRNAs in the cellular response against the virus by using transcriptomic data obtained from nasopharyngeal swabs in a cohort of patients with different SARS-CoV-2 viral loads. We determined the existence of a transcriptomic dysregulation pattern in SARS-CoV-2 infected patients that is mainly represented by upregulated genes. In those patients with higher viral loads, a significant proportion of the upregulated transcriptome is composed by lncRNA transcripts.

To decipher the possible role of the upregulated lncRNAs during SARS-CoV-2 infection, we used a systems biology-based approach. In absence of biological validation, the “guilty by association” principle can be applied to predict the function of a group of lncRNAs [73, 74]. Using the ncFANS platform, the embedded NET algorithm and a subsequent enrichment for GO-terms and metabolic pathways, we revealed a striking molecular fingerprint of the differentially upregulated lncRNAs that pointed to their functional correlation with lymphocyte activation and cytokine signaling (Fig. 2a and 2b). The intertwined relationship between cytokine signaling and lncRNA regulation provides a feedforward/feedback regulatory mechanism in the control of cellular responses to cytokines [75]. Here, we analyzed the molecular signature of the upregulated lncRNAs during SARS-CoV-2 infection (Fig. 2c), determining their involvement in the regulation interferon-regulated inflammatory response.

Viral-induced lncRNA upregulation has been observed in respiratory viruses such as SARS-CoV [76], influenza [77], and SARS-CoV-2 [78, 79], using cellular or animal models on infection. Notably, in primary normal human bronchial epithelial cells (NHBE) infected with SARS-CoV-2, the transcriptomic analysis revealed the overexpression of interferon-responsive genes, namely IRF9, IFIT1, IFIT2, IFIT3, IFITM1, MX1, OAS2, OAS3, IFI44 and IFI44L, together with the induction of an acute inflammatory response and activation of tumor necrosis factor (TNF) [79]. In this model, the interferon response was also linked to the overexpression of at least 18 different lncRNAs. However, the lncRNAs induced in NHBE cells after viral infection are distinct from the signature we determined in the analyzed human nasopharyngeal samples, probably due to the cell and tissue specificity of lncRNA expression [79]. Involvement of interferon-responsive lncRNAs in the regulation of viral infections is a widespread phenomenon where lncRNAs usually have a detrimental effect over viral infections. For instance, the interferon-stimulated lncRNA (ISR) is actively induced after influenza virus infection in animal models and it is involved in the control of viral replication [15]. Interestingly, interferon-independent lncRNAs are frequently hijacked by viruses like influenza to promote their replication using different molecular mechanisms that involve the stabilization of the viral genome and its replication machinery [77]. In particular, Epstein Barr Virus is known to utilize IncRNAs in both lytic and latent phase infections [80] providing further evidence for the idea that many of these viral systems may be hijacking the same mechanism through LncRNAs. For EBV this is particularly interesting because of the role that EBV seems to play in long Covid [81], and how SARS-CoV-2 infection can induce latent phase activation of EBV itself.

The lncRNAs components of the non-coding transcriptional signature induced in nasopharyngeal samples during SARS-CoV-2 infection, NRIR, LINC02068, BISPR, USP30-AS1, MIR155HG and AL512306.2, are significantly upregulated (Fig. 3). Interestingly, NRIR, BISPR, USP30-AS1 and AL512306.2 lncRNAs were also detectable at higher levels in blood samples of SARS-CoV-2 patients. NRIR, BISPR and MIR155HG levels in nasopharyngeal samples are correlated with the viral load, quantified as the expression of the S-protein coding gene (Fig. 4). LncRNA NRIR, formerly designated as lncRNA-CMPK2, was first characterized as an interferon-responsive transcript that exerts a negative regulatory effect over the interferon defensive pathway. Using in vitro models of hepatitis C virus (HCV) infection, the knockdown of NRIR gene resulted in a marked reduction in HCV replication in interferon-stimulated hepatocytes, suggesting that it could affect the antiviral role of interferon [82]. In other viral infections such the Crimean-Congo hemorrhagic fever, NRIR has been also shown to be upregulated [68]. The molecular partners associated with NRIR regulatory action are still not characterized, however preliminary evidence suggested that NRIR could act via recruitment of chromatin-remodeling enzymes [51, 82]. In contrast, BISPR lncRNA is a divergent non-coding transcript generated from the promoter of the BST2 gene, that acts as a transcriptional enhancer of the BST2 gene [83]. BST2 protein, also known as Tetherin, is an interferon-induced transmembrane protein that has been involved in the inhibition of the replication of RNA viruses by controlling the release of the viral particles [84, 85] or by inducing the apoptosis of the host cells [86]. MIR155HG lncRNA has been also characterized as a positive regulator of the cellular immune response in influenza infections. In cellular models, MIR155HG showed an inhibitory effect on the expression of protein tyrosine phosphatase 1B (PTP1B) during the infection with influenza A virus, that could be directly related with the increase in the expression of interferon-beta [53]. Taken together, our data showed that the lncRNA signature induced by SARS-CoV-2 infection in the host includes counteracting regulators of cellular immunity: NRIR that could act as a promoter of viral replication by a negative regulation of interferon response, and BISPR and MIR155HG, that could act as antiviral lncRNAs. Understanding the balance between promoters and inhibitors of the viral progression will be essential to derive better effective therapeutics specially for severe cases of infection.

We performed additional analysis to infer the molecular consequences that might be associated with the lncRNA signature induced by SARS-CoV-2 infection. Considering that the regulatory lncRNA action is exerted by their contribution to the stability and function of macromolecular complexes, we applied a functional association method based on the IHNLncSim algorithm [44]. This strategy allowed us to characterize additional lncRNAs that could be participating in the host response to SARS-CoV-2 infection. Moreover, we combined protein-lncRNA interaction data to give insights about the possible involvement of the lncRNA-RBPs axis in the overall cellular response against infection. Globally, this functional analysis of the experimentally validated events involving RBPs and the virus-induced lncRNA signature showed a striking pattern related with the regulation of mRNA metabolism, stability, and processing by splicing (Fig. 5a and 5b). Such connection between viral infections, lncRNAs and splicing has been demonstrated for other RNA viruses. Using Zika-infected human neural progenitor cells, Hu and coworkers observed that the transcriptional lncRNA shift induced by the virus was accompanied by a specific pattern of splicing events that affected genes involved in cell proliferation, apoptosis, and differentiation [87]. Similarly, proteomics and transcriptomic data obtained in cellular models of SARS-CoV-2 infection determined that the viral NSP16 protein binds to the mRNA recognition domains of the U1 and U2 spliceosomal RNAs, suppressing the canonical splicing events [88].

Among the viral-induced lncRNAs, we selected a specific signature composed by six elements (NRIR, BISPR, MIR155HG, USP30-AS1, FMR1-IT1, and U62317.2), for functional association studies [44]. Except for BISPR lncRNA, the functional lncRNA signature is composed of lncRNAs with low-density of RBP-binding sites, suggesting their involvement in specific regulatory events instead of being involved in the capture of RBPs by sponging (Fig. 5c). Despite the previous functional characterization of some of the lncRNAs induced by SARS-CoV-2 infection [89-91], this regulatory network is not complete, and the presence of the viral RNA genome cannot be neglected. To support this idea, Schmidt and coworkers recently applied an antisense-based purification protocol coupled to mass spectrometry to determine the protein host-cell interactome in SARS-CoV-2 infection models [32]. The results presented a detailed RBP-virus interactome, showing how the external modulation of the levels of specific host proteins such LARP1 or CNBP could be used as therapeutic targets to restrict the viral replication [32].

The rationale of our proposed model is based on the integration of RBP-lncRNA interaction data combined with an additional layer of complexity established by the presence of the viral RNA genome. Analyzing the validated protein interactome of the lncRNA signature induced by SARS-CoV-2 infection and the proteomic data of the host-interacting proteins, we were able to determine the presence of RBPs that are simultaneously interacting with the SARS-CoV-2 genome and with the overexpressed lncRNAs (Fig. 6a). This group of RBPs comprises MSI2, IGF2BP1, IGF2BP2, UPF1, HNRNPA1, DDX3X, PCBP2 and LIN28B proteins.

Except for PCBP2, UPF1 and LIN28B proteins, the RBPs able to interact with the viral genome and the upregulated lncRNAs, followed an expression time course that also accompanied the SARS-CoV-2 infection (Fig. 6b) [25, 92]. This evidence suggested that the competition between host lncRNAs and the viral genome for the binding of specific RBPs could control the cellular response and the viral replicative cycle [93]. Our working hypothesis is supported by research on the roles of selected RBPs in other viral infections. For instance, UPF1, a highly processive helicase required for non-sense mediated decay (NMD) [94], has been described as an essential factor for the completion of the replicative cycle of HIV [95]. Moreover, DDX3X, another RNA helicase, could be probably the most relevant element of this lncRNA-RBP interaction network. In hepatitis B virus, DDX3X helicase is an essential factor since it can restrict the replicative cycle of the virus at the transcriptional level [96]. Additional evidence obtained in influenza virus infections also characterize DDX3X helicase as an essential and driving factor for the host cell response against influenza A virus (IAV) infection [97]. DDX3X is critical to orchestrate a multilayer antiviral innate response during infection, coordinating the activation of the inflammasome, assembly of stress granules, and type I interferon (IFN) responses. The loss of DDX3X expression in myeloid cells caused an increase of susceptibility to pulmonary infections and morbidity in IAV-infected mice [97]. However, the roles of DDX3X helicase as a promoting or preventing factor for viral infections appear to be virus-specific, as exemplified by the comparison of hepatitis B and C viruses [98]. In SARS-CoV-2 infection, the DDX3X protein levels are upregulated during infection (Fig. 6b) suggesting its involvement in the cellular response against the virus. Also, among RBPs able to crosstalk with the viral genome and the upregulated lncRNAs is IGF2BP2, a multifaceted RBP able to control multiple metabolic processes [99]. There is no current evidence of a direct role of IGF2BP2 in the response against viral infections, but recent data connected its function to the lncRNAs. In thyroid cancer, IGF2BP2 is an oncogenic factor since it enhances the expression of the HAGLR lncRNA and the cellular proliferation [100]. Also, in non-small-cell lung cancer, IGF2BP2 enhances the proliferation of tumor cells by binding to the oncogenic MALAT1 lncRNA and increasing its stability [101]. In our model, IGF2BP2 can interact with BISPR and MIR155HG lncRNAs, two of the core elements of the functional signature induced by SARS-CoV-2 infection that are involved in the host cell innate immune response.

In summary, SARS-CoV-2 infection induces an upregulation of specific lncRNAs that are functionally correlated and involved in the innate immune response. These lncRNAs form part of an intricate network of interactions that involve specific RBPs, represented by helicases such as UPF1 and DDX3X, both regulators of RNA stability and surveillance. The presence of the viral RNA(+) genome introduces another layer of complexity since it can compete with lncRNA for the binding of the RBPs, being possibly involved in the modulation of the cellular response against infection and the viral progression. The specific functions of these interactions would require further biological validation and could open new possibilities for understanding the host components required for SARS-CoV-2 progression and designing new targeted antiviral therapeutics.

## 5. ABBREVIATIONS

IAV: Influenza A virus
HIV: human immunodeficiency virus
lncRNA: long non-coding RNA
mRNA: messenger RNA
RBP: RNA-binding protein
RNA: ribonucleic acid

## 6. ACKNOWLEDGEMENTS

The authors would like to thank Francisco Enguita Jr. for his valuable insight and help in the design and conceptualization of the manuscript.

## 7. COMPETING INTERESTS

The authors have declared that no competing interest exists.

## Notes

### Competing Interest Statement

The authors have declared no competing interest.

